# F-BAR proteins CIP4 and FBP17 function in cortical neuron radial migration and process outgrowth

**DOI:** 10.1101/2024.10.25.620310

**Authors:** Lauren A. English, Russell J. Taylor, Connor J. Cameron, Emily A. Broker, Erik W. Dent

## Abstract

Neurite initiation from newly born neurons is a critical step in neuronal differentiation and migration. Neuronal migration in the developing cortex is accompanied by dynamic extension and retraction of neurites as neurons progress through bipolar and multipolar states. However, there is a relative lack of understanding regarding how the dynamic extension and retraction of neurites is regulated during neuronal migration. In recent work we have shown that CIP4, a member of the F-BAR family of membrane bending proteins, inhibits cortical neurite formation in culture, while family member FBP17 induces premature neurite outgrowth. These results beg the question of how CIP4 and FBP17 function in radial neuron migration and differentiation *in vivo*, including the timing and manner of neurite extension and retraction. Indeed, the regulation of neurite outgrowth is essential for the transitions between bipolar and multipolar states during radial migration. To examine the effects of modulating expression of CIP4 and FBP17 *in vivo*, we used *in utero* electroporation, in combination with our published Double UP technique, to compare knockdown or overexpression cells with control cells within the same mouse tissue of either sex. We show that either knockdown or overexpression of CIP4 and FBP17 results in the marked disruption of radial neuron migration by modulating neuronal morphology and neurite outgrowth, consistent with our findings in culture. Our results demonstrate that the F-BAR proteins CIP4 and FBP17 are essential for proper radial migration in the developing cortex and thus play a key role in cortical development.

**SIGNIFICANCE STATEMENT:** During embryonic development, radial migration of newly born cortical neurons is a complex process that underlies the proper formation of the neocortex, the outermost layers of neurons in the brain. Disruptions in radial migration results in profound effects on cognitive function and can lead to devastating developmental disabilities. To better understand this critical process in brain development we examined two members of the F-BAR family of membrane bending proteins, CIP4 and FBP17, which are present in the developing brain. We demonstrate that intracellular concentrations of these proteins must be tightly regulated. Increasing or decreasing levels of either protein has profound effects on neuronal morphology and proper radial migration, suggesting they are key players in cortical development.

## INTRODUCTION

The development of the mouse cerebral cortex requires precisely timed birth, migration, and maturation of millions of neurons. Pyramidal excitatory neurons are born in the ventricular zone, adjacent to the lateral ventricles (Parnavelas, 2000), then undergo a series of stereotyped morphological changes as they migrate radially from the ventricular zone to the cortical plate, taking their positions in a newly forming six-layer cortex (Noctor et al., 2004). Newly born pyramidal neurons start out as spherical cells that extend short neurites termed leading and trailing processes. They begin their journey as a bipolar neuron in the ventricular zone, but in the subventricular zone they pause and develop a multipolar morphology. Subsequently, they retract these extraneous processes, at which point they resume radial migration as a bipolar neuron, until they reach their destination in the cortical plate, where they extend an axon toward the ventricular zone and a dendrite toward the pial surface (Noctor et al., 2004).

Rho GTPases, and their associated GEFs and GAPs, are well known to play important roles in radial migration of cortical neurons (Govek et al., 2010; Azzarelli et al., 2014; Xu et al., 2019). Activated Rho GTPases, including Rac1, Cdc42 and RhoA, are recruited to the plasma membrane and interact with selected membrane-associated proteins (Bement et al., 2024). One family of proteins that plays important roles in Rho GTPase signaling cascades are the F-BAR family of proteins. Although certain F-BAR proteins play key roles in cortical migration and neuronal development (Guerrier et al., 2009; Charrier et al., 2012), the roles of other F-BAR family members have been little studied, including the Cdc42 interacting protein 4 (CIP4) family of proteins.

The CIP4 family proteins are a subgroup of F-BAR proteins that are expressed in many tissues and act as polymeric, membrane binding complexes that interact with activated Rho GTPases and recruit actin associated proteins (Aspenstrom, 2009; Snider et al., 2021). The CIP4 subfamily is comprised of three members: CIP4, Formin binding protein 17 (FBP17) and Transducer of Cdc42-dependent actin assembly protein 1 (TOCA1) (Aspenstrom, 2009). These proteins typically sense and induce membrane curvature during endocytosis (Itoh et al., 2005; Henne et al., 2007). However, we have shown that in dissociated cortical neurons CIP4 does not appear to function in membrane tubulation, but instead localizes to the protruding, peripheral plasma membrane (Saengsawang et al., 2012; Saengsawang et al., 2013; Taylor et al., 2019). Sustained overexpression of CIP4 inhibits neurite formation in dissociated cortical neurons, while newly plated neurons from CIP4 knockout mice initiate neurites prematurely (Saengsawang et al., 2012). CIP4 is expressed in the early embryonic cortex, with levels decreasing until birth, at which point it remains largely absent (Saengsawang et al., 2012). Curiously, FBP17 has an opposite effect in neurons compared to CIP4. FBP17 functions in endocytosis by localizing to discrete tubules in the cytoplasm and overexpression of FBP17 induces premature and excess neurite outgrowth (Taylor et al., 2019). Moreover, unlike CIP4, brain-wide FBP17 expression increases in late embryonic development and peaks in adulthood (Wakita et al., 2011). These findings suggest that CIP4 and FBP17 may have important and distinct roles in embryonic cortical neuron development.

To investigate the roles of CIP4 and FBP17 in radial migration we utilized *in utero* electroporation (IUE) to introduce plasmid DNA into newly born neurons along the lateral ventricles in the embryonic mouse cortex (Saito and Nakatsuji, 2001; Tabata and Nakajima, 2001). Recently, we increased the rigor and reproducibility of IUE through a technique we describe as Double UP (Taylor et al., 2020). Double UP generates green cells containing no manipulation, and magenta cells that express a manipulation of interest (overexpression or knockdown) in the same electroporated region of cortex. This technique allows direct comparisons of control and manipulated neurons within the same section of cortex, as neurons dynamically migrate and change their morphology. Applying this technique, we show that CIP4 and FBP17 play critical, distinct roles in radial migration.

## METHODS

### DNA Constructs

Double UP mNeon to mScarlet is available through Addgene (#125134). pCAG-iCre was a gift from Wilson Wong (Addgene plasmid #89573). pSico PGK Puro was a gift from Tyler Jacks (Addgene plasmid #11586). The two shRNA sequences targeting mouse CIP4 are: (1) 5’- GTCTGGAGCTGGCTAAGTA-3’ and (2) 5’-GTCTGGAGCTGGCTAAGTA-3’. The two shRNA sequences targeting mouse FBP17 are: (1) 5’-GCAAGCTCTGGCCATTCAT-3’ and (2) 5’- GTCGTAGAAGCCTATAAGT-3’. The shRNA sequence for the scramble construct is 5’- GGCGGAGATTTCTGACTGA-3’. Human CIP4 (short isoform – 545 aa, NM_004240.4) and human FBP17 (long isoform – 617 aa, NM_0.15033.3) were utilized for these experiments (Taylor et al., 2019). Human CIP4 short and FBP17 long isoforms that are C-terminally tagged with either EGFP or mScarlet are available from Addgene (#178363-178366). Further information and requests for resources and reagents should be directed to and will be fulfilled by the lead contact, Erik Dent.

### Animal Models

All mouse procedures were approved by the University of Wisconsin Committee on Animal Care and were in accordance with NIH guidelines. Timed matings of Swiss Webster mice (Inotiv) were performed, with the morning of sperm plug visualization considered E0.5. IUE was performed at E14.5, with embryos perfused either two or four days later, as specified in the text. Both male and female embryos were used, with the gender of embryos not recorded. Pregnant females were housed individually. Prior to becoming pregnant, females were housed with 3-4 other females.

### In Utero Electroporation (IUE)

Before injection, 280ng of pCAG-iCre was mixed with 30µg of Double UP, Double UP CIP4, or Double UP FBP17. For knockdown studies, 280ng pCAG-iCre was mixed with 15µg Double UP and 30µg of either pSico-CIP4, pSico-FBP17, or pSico-scramble. Plasmid DNA was then combined with Fast Green FCF to a final concentration of 0.05% and loaded into pulled capillary needles. The dam was anesthetized with isoflurane and a laparotomy was performed, exposing the embryos. The embryos were gently removed from the abdominal cavity. Capillary needles were inserted into the lateral ventricles, and approximately 0.25-0.5µL DNA/Fast Green FCF was injected using a PicoSpritzer II (Parker Instrumentation). Electrical current was passed across the head, in five pulses of 40 volts each lasting 100ms on and 900ms off, with a CUY21 Electroporator (Bex Co. LTD). After the last embryo was electroporated, the embryos were inserted back into the mother, and the laparotomy was sutured closed. Dams were given 20 mg/kg of Carprofen and 3.25 mg/kg extended-release buprenorphine (Ethiqa) S.C. Dams were monitored for signs of pain every twelve hours for two days. Embryos were allowed to develop normally for 2-4 days, as indicated.

### Generation of CIP4-mScarlet Transgenic Mouse

A cassette coding for Linker-LoxP-3xHA-Stop Codon-LoxP-mScarlet was cloned using Gibson Assembly. Genomic DNA was isolated from wild type mouse liver and used to clone a 1.5kb 5’ homology arm, and a 2.5kb 3’ homology arm located immediately around the stop codon of CIP4. These three components were combined using Gibson Assembly, and the resulting plasmid DNA was used as a template for Crispr/Cas9. A guide RNA (GAACCCCACCAGAGGGGGACG(AGG)) was validated to cut genomic DNA, and then inserted with Cas9 protein and linearized homology plasmid into fertilized mouse oocytes. Animals were screened for presence or absence of a transgene via PCR, and a female was found to be mosaic for the insert. The transgene and surrounding 1KB of genomic DNA were sequence verified. The female was used as the founder of the colony. Once sufficient animals were positive for the transgene, a heterozygous female was crossed to a ß-Actin Cre Mouse (Jax 019099), to permanently recombine the transgene to code for CIP4-Linker-LoxP-mScarlet. Both lines (CIP4- Linker-LoxP-mScarlet and CIP4-Linker-LoxP-3xHA-Stop Codon-LoxP-mScarlet) are being maintained by the Dent Lab and are available upon request. Genotyping is performed using ACCAGGGGATGTAGCAGTTG (Forward) and CACGTGGGCAGGAATAAAGT (Reverse) primers.

### Tissue Collection and Sectioning

For both IUE and endogenous CIP4-mScarlet/ WT experiments, embryos were exposed via laparotomy after deep anesthesia of the pregnant dam with isoflurane. Embryos were removed from the uterus one by one and perfused by opening the chest cavity, making a small incision in the right atrium and inserting of a 25-gauge needle into the left ventricle. Through this needle, the animal was perfused with approximately 1mL of sterile saline and 3mL of 4% paraformaldehyde in PBS (PFA) at the rate of approximately 1.25mL per minute with a perfusion pump (Instech). Following perfusion, heads were removed and placed in 4% PFA at 4°C overnight. After the last embryo was perfused, the dam was euthanized via live decapitation. P10 animals for the endogenous CIP4-mScarlet/WT experiments were given 120 mg/kg of sodium pentobarbital before intracardial perfusion as described above.

After 16 hours in 4% PFA at 4°C, heads were transferred to PBS and brains were removed from skulls. The brains were placed in 6% low melt agarose and allowed to set on ice. After the agarose hardened, the brains or heads (only Fig. 5 E12.5 and E14.5) were sectioned on a Leica VT1000S vibratome at 100µm in phosphate-buffered saline (PBS). Sections from animals that underwent IUE were stored for less than one week in PBS before being stained with 4′,6- diamidino-2-phenylindole (DAPI) and imaged. Sections were incubated in DAPI (2.4nM in 0.4% Triton/PBS) for 1 hour with gentle rotation at room temperature. Slices were washed in PBS and mounted onto slides with Aqua-Poly Mount (Polysciences). Slides were allowed to dry for at least one hour and then imaged within two days. Sections from endogenous CIP4-mScarlet/WT experiments were incubated in primary antibody (anti-mScarlet 1:500, Rockland) diluted in blocking solution (10% Normal Goat Serum-NGS (Innovative), 0.4% Triton (Sigma), 2% Bovine Serum Albumin (Sigma), 1% Glycine (Sigma) in PBS) overnight at 4°C. The following day, the primary antibody was washed off with 10% NGS 3 times and a secondary antibody was applied in addition to DAPI for 1 hour, gently rotating at room temperature. Sections were mounted as described previously.

### Imaging, Data Collection and Quantification

Imaging was performed on a Zeiss LSM 800 confocal microscope. For migration analysis **(Fig. 1- 4)**, 12 optical sections were obtained, each 1µm apart. 2x2 tiles were collected with a 20x/0.8NA Plan Apochromat objective, with 2x averaging. For fixed morphology analysis **(Fig. 3, 4)**, 80-120 optical sections were obtained through the entirety of the 100µm section, each at 0.5µm. Although the entire section was imaged in the z-plane, less than 100µm was needed due to tissue compression. These images were also collected with a 20x/0.8NA Plan Apochromat objective, with 2x averaging. For IUE experiments, gain/laser power were altered between each image set to optimize image quality. Tiles were stitched together using the stitching tool in Zen 2.3 (Zeiss) software. For imaging of endogenous CIP4-mScarlet **(Fig. 5)**, 58 optical sections were obtained, each 1µm apart. Images were collected with a 20x/0.8NA Plan Apochromat objective, with 4x averaging, and presented as maximum projections of the six brightest continuous sections.

**Figure 1.**
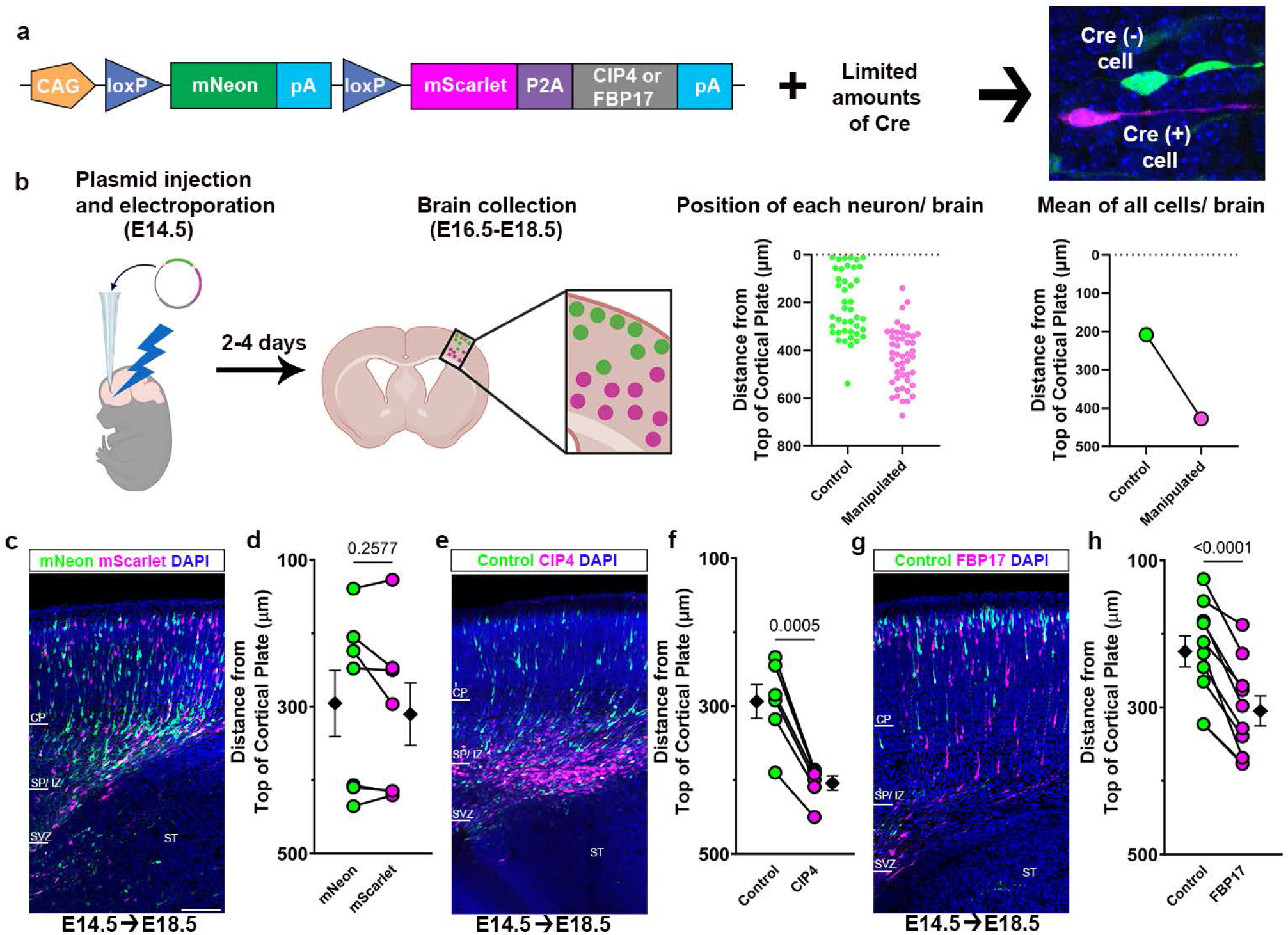
Overexpression of CIP4 or FBP17 decreases migration at E14.5+4 days. **(a)** Representation of the Double UP overexpression plasmid construct used in these experiments. Neurons that are transfected with only the Double UP plasmid express mNeon- green (Cre(-) cell). When limited amounts of Cre are co-transfected into about half the neurons, these neurons express mScarlet (mNeon-green is excised) and unlabeled CIP4 or FBP17 (Cre(+) cell). **(b)** Schematic of experimental design and quantitation procedure. **(c, e, g)** Representative images of empty Double UP (c), Double UP + CIP4 (e), and Double UP + FBP17 (g) four days after electroporation at E14.5 (E14.5◊E18.5). CP= cortical plate, SP/IZ= subplate/intermediate zone, SVZ= subventricular zone, ST= striatum, V= lateral ventricle. Scale bar 100µm. **(d, f, h)** Dot plots of migration of green and magenta cells. Each dot represents cumulative mean distance from the top of the cortical plate for all labeled neurons in a single coronal brain section. Connected dots indicate measurements were made in the same coronal section. Black diamonds and bars represent cumulative mean ± SEM. Paired t-test (two-tailed). (d) 7 brains, mNeon: mean 295±45.0µm, 1316 cells, mScarlet: 310±44.4µm, 1055 cells. (f) 6 brains, Control: 294±23.0µm, 745 cells, CIP4: 404±9.8µm, 731 cells. (h) 10 brains, Control: 224±21.0µm, 2144 cells, FBP17: 305±20.5µm, 1828 cells. (b) Created in BioRender. English, L. (2024) BioRender.com/u35d157.

Images of cortical neuron radial migration were analyzed and migration distances were calculated using the TRON software program (Taylor et al., 2020). Process number and length of E14.5 +2 neurons were calculated in Zen 2.3 (Zeiss) software. Neuronal polarity was calculated from process number data. All experimenters were blinded to condition.

### Western Blot Analysis

For validation of shRNA constructs **(Fig. 2d, e, j, k)** HEK293 cells were transfected with a post-cre recombined version of the pSico plasmid and either mouse CIP4-mScarlet **(Fig. 2d, e)** or mouse FBP17-mScarlet **(Fig. 2j, k)** using Lipofectamine 3000 (Invitrogen) following the manufacturers’ protocol. After 48 hours, cells were washed once with cold PBS before being lysed using 300μL NP-40 Lysis Buffer (Invitrogen) with Complete Mini (Roche). Lysate was spun at 21,000*g* for 10 minutes, and supernatants were flash-frozen and stored at -80°C until use. Samples were thawed and loaded onto a 4–10% SDS-Page gel, then transferred to PVDF membrane (Millipore). Membranes were blocked in 5% milk in 0.1% TBS-T, incubated with primary antibody overnight at 4°C, and blotted with an HRP-conjugated secondary antibody for 1 hour at RT. Antibodies used were mouse anti-CIP4 (1:1000, BD Biosciences), Rabbit anti-FBP17 (1:500, Proteintech), goat-anti-mouse HRP (1:10000, Jackson), and goat-anti-rabbit HRP (1:5,000, Thermo Scientific). Protein bands were visualized using Pierce ECL Western blotting substrate (Thermo Scientific).

**Figure 2.**
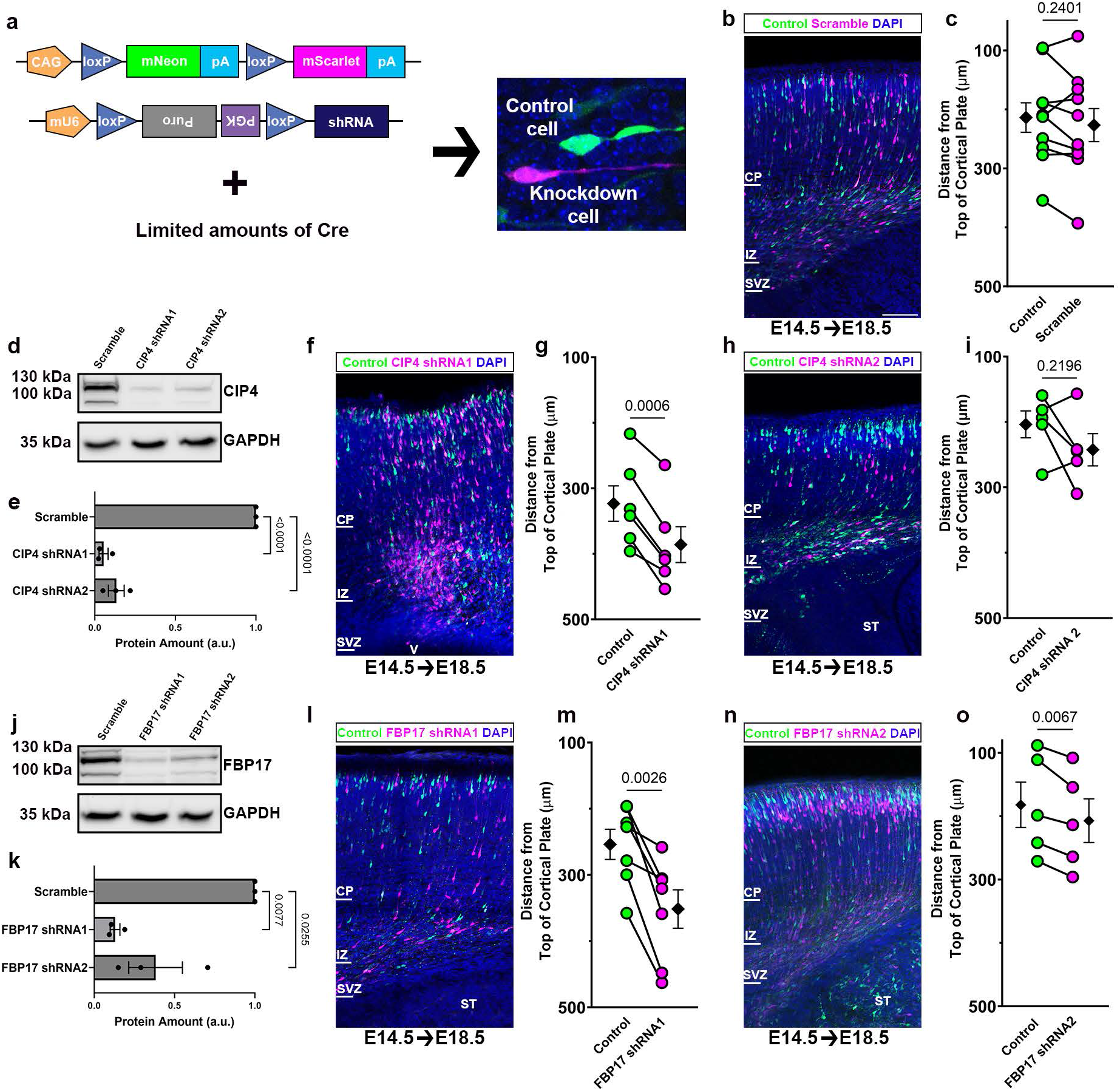
Knockdown of CIP4 or FBP17 decreases migration at E14.5+4 days. **(a)** Representation of the Double UP and pSico plasmid constructs used in these experiments. Neurons that are transfected with both the Double UP plasmid and a floxed, inverted PGK- puromycin cassette express only mNeon-green and no shRNA (Control cell). When limited amounts of Cre are co-transfected into about half the population of neurons, these neurons express mScarlet (mNeon is excised) and shRNA (PGK-puro cassette excised) targeting either CIP4 or FBP17 (Knockdown cell). **(b)** Representative image of Double UP + pSico scramble shRNA four days after electroporation at E14.5 (E14.5◊E18.5). Scale bar 100µm. **(c)** Dot plot of migration of green and magenta cells. Each dot represents cumulative mean distance from the top of the cortical plate for all cortical neurons in a coronal brain section. Connected dots indicate measurements were made in the same coronal section. **(d)** Western blot of HEK293 cells transfected with mouse CIP4 and post-Cre pSico plasmids expressing either scramble or one of two CIP4-targeted shRNAs. **(e)** Quantification of western blots in (d). **(f, h)** Representative images of Double UP + pSico CIP4 shRNA1 (f) and shRNA2 (h) four days after electroporation at E14.5 (E14.5◊E18.5). **(g, i)** Dot plots of migration of green and magenta cells for CIP4 shRNA1 (g) and shRNA2 (i). **(j)** Western blot of HEK293 cells transfected with mouse FBP17 and post-Cre pSico plasmids expressing either scramble or one of two FBP17-targeted shRNAs. **(k)** Quantification of western blots (j). **(l, n)** Representative image of Double UP + pSico FBP17 shRNA1 (l) and shRNA2 (n) four days after electroporation at E14.5 (E14.5◊E18.5). **(m, o)** Dot plots of migration of green and magenta cells for FBP17 shRNA1 (m) and shRNA2 (o). Black diamonds and bars represent cumulative mean ± SEM. Paired t-test (two-tailed). (c) 10 brains, Control: 214±25.0µm, 2453 cells, scramble: 227±28.0µm, 2142 cells. (g) 6 brains, Control: 324±27.0µm, 1669 cells, CIP4 shRNA1: 386±27.4µm, 1631 cells. (i) 5 brains, Control: 204±20.5µm, 921 cells, CIP4 shRNA2: 243±24.7µm, 798 cells. (m) 7 brains, Control: 254±22.8µm, 959 cells, FBP17 shRNA1: 352±29.1µm, 1095 cells. (o) 5 brains, Control: 182±35.8µm, 1269 cells, FBP17 shRNA2: 207±34.5µm, 1082 cells.

### Quantification and Statistical Analysis

Radial migration distances, number of processes, and length of longest process data were averaged for each color in each brain. Paired, two-tailed, t-tests were performed on the averages from each brain. Neuronal polarity was analyzed by a two-way ANOVA with multiple comparisons. Complete data is available upon request. All statistical tests were performed in Prism 8 (GraphPad). Several graphs are presented as Superplots (Lord et al., 2020; Goedhart, 2021).

## RESULTS

### Overexpression of either CIP4 or FBP17 results in migration defects

Radial migration relies on a stereotyped series of extensions and retractions of minor processes during migration from the ventricular zone to the cortical plate (Noctor et al., 2004). Sustained CIP4 or FBP17 overexpression markedly inhibits or promotes the formation and growth of neurites in cortical neurons *in vitro,* respectively (Saengsawang et al., 2012; Saengsawang et al., 2013; Taylor et al., 2019). Since CIP4 expression decreases throughout embryonic cortical development (Saengsawang et al., 2012), we hypothesized that maintaining neuronal CIP4 expression *in vivo* would result in neurons lacking processes and negatively affect radial migration. Additionally, we hypothesized that sustained FBP17 overexpression *in vivo* would result in neurons with excess processes and likewise lead to a migration defect. To test these hypotheses, we utilized *in utero* electroporation at embryonic day 14.5 (E14.5) in combination with the Double UP technique developed in the lab (Taylor et al., 2020). Double UP allowed us to generate both control (mNeon-green) and manipulated (mScarlet, shown in magenta) cells in the same brain by utilizing limiting amounts of Cre to recombine plasmids in roughly half of the transfected cells **(Fig. 1a, b)**. This technique eliminates the need for section matching, decreasing variability between conditions (Taylor et al., 2020). Double UP also allows for the statistical comparison of matching mean migration distances in control and manipulated cells **(Fig. 1b)**. For these experiments we created two versions of the Double UP construct to either overexpress mScarlet + unlabeled CIP4 (Double UP-CIP4) or mScarlet + unlabeled FBP17 (Double UP-FBP17) **(Fig. 1a, b)**, while control neurons expressed only mNeon.

We first confirmed our previous work (Taylor et al., 2020) showing that expression of neither mNeon nor mScarlet fluorescent proteins alone differentially affect neuronal migration **(Fig. 1c, d)**. Note that although the average distance migrated varied between experiments, both green and red cells, within the same brain section, generally migrated similar distances. Proceeding to our experimental conditions, we found that CIP4 overexpression in cortical neurons (magenta) from E14.5 to E18.5 (E14.5 +4) resulted in a consistent, robust reduction of migration compared to green controls **(Fig. 1e, f)**. The majority of CIP4 overexpressing cells remained in the ventricular and subventricular zone **(Fig. 1e)**. FBP17 overexpression for four days (E14.5 +4) showed a consistent, marked reduction in migration as well **(Fig. 1g, h)**. The mScarlet/FBP17 positive cells were primarily located in the intermediate zone compared to controls **(Fig. 1g)**. These results suggest that overexpression of either CIP4 or FBP17 markedly inhibits radial migration of cortical neurons.

### Knockdown of CIP4 or FBP17 decreases migration in the intermediate zone at E14.5 +4

Overexpression of either CIP4 or FBP17 resulted in dramatic migration defects. However, it is important to consider that overexpression of any protein may induce cellular defects from the inherent disruption of endogenous protein level. Therefore, we elected to examine whether decreasing expression of CIP4 and FBP17 would be similarly disruptive to radial migration. Utilizing Double UP in combination with a Cre-dependent shRNA vector, “pSico” (Ventura et al., 2004), we designed shRNAs to knockdown CIP4 and FBP17 *in utero*, while maintaining control (green) and knockdown (magenta) cells in the same slice **(Fig. 2a)**. We first tested a scrambled shRNA sequence as a control, electroporating Double UP combined with a scrambled shRNA. This control experiment showed equivalent levels of migration between magenta and green cells **(Fig. 2b, c)**.

We utilized two different shRNAs for both CIP4 and FBP17 and tested their effectiveness in HEK293 cells overexpressing either mouse CIP4 (mCIP4) or mouse FBP17 (mFBP17). Quantification of western blots of HEK293 lysates showed significant knockdown of both expressed mCIP4 **(Fig. 2d, e)** and mFBP17 **(Fig. 2j, k)**. Since CIP4 protein expression normally decreases substantially during prenatal cortical development (Saengsawang et al., 2012), we were unsure if we could knock down the protein to levels that would result in a migration phenotype. However, we discovered that CIP4 knockdown with the stronger shRNA1 resulted in a robust reduction of migration **(Fig. 2d-g)**, while the weaker shRNA2 resulted in an inconsistent migration defect **(Fig. 2d, e, h, i)**. Cells expressing CIP4 shRNA1 consistently clustered in the intermediate zone below control cells **(Fig. 2f)**. CIP4 shRNA2 showed inconsistent results with cells more spread throughout the intermediate zone and cortical plate **(Fig. 2h)**.

We then tested whether our FBP17 shRNAs resulted in defects in radial migration. Both shRNAs induced knockdown of FBP17 that resulted in decreased radial migration **(Fig. 2l-o)**. FBP17 shRNA1 halted migration in the lower cortical plate and intermediate zone **(Fig. 2l)**, while FBP17 shRNA2 expressing cells migrated into the cortical plate but not as far as controls **(Fig. 2n)**. Together, these results suggest that either increasing or decreasing expression of CIP4 or FBP17 consistently and markedly disrupts migration of radially migrating cortical neurons.

### Overexpression of CIP4, but not FBP17, negatively affects migration by inhibiting process initiation at E14.5 + 2 days

We next examined an earlier timepoint after electroporation (E15.4 +2) to glean insight into the mechanism underlying the disruption of radial migration induced by varying levels of CIP4 and FBP17 expression. Migrating neurons assume a variety of morphologies during their migration from the ventricular zone to the cortical plate, transitioning between bipolar and multipolar shapes, corresponding to the zone of cortex they inhabit (Noctor et al., 2004). Examination of neurons two days after electroporation allowed for the direct comparison of neuronal morphology between control and manipulated neurons within a similar region of cortex, because most cells had not yet migrated far from the ventricular zone. Again, to confirm that our findings were due to CIP4 or FBP17 overexpression, we utilized Double UP, which induces expression of mNeon or mScarlet after electroporation. Green and magenta cells showed no difference in migration **(Fig. 3a, b).** We also compared the morphology of cells expressing Double UP at E14.5 +2. Comparison of morphologies at this stage of development was more appropriate than at E14.5 +4, when migrating neurons are in different cortical zones and take on a different morphology corresponding to their location in cortex. Green and magenta cells had similar morphologies, with no difference in the number or length of processes **(Fig. 3c-f)**. Additionally, we analyzed the polarity of the migrating cells and found most cells exhibited a multipolar morphology at E14.5+2 in both Double UP conditions **(Fig. 3g)**.

**Figure 3.**
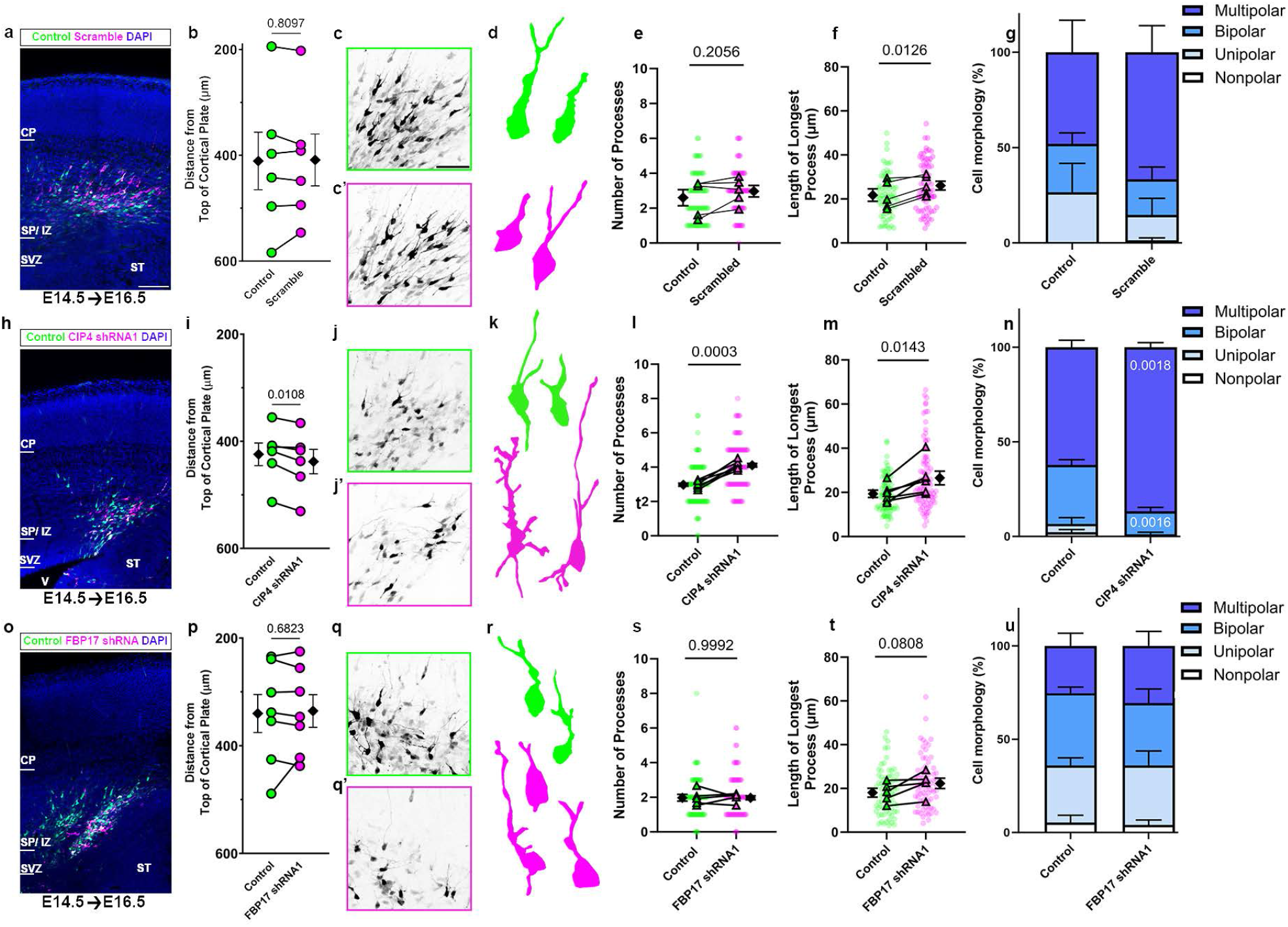
CIP4, but not FBP17, overexpression inhibits migration by decreasing number and length of processes. **(a, h, o)** Representative images of migration after electroporation with Double UP (a) Double UP CIP4 (h) and Double UP FBP17 (o) two days after electroporation at E14.5 (E14.5◊E16.5). CP= cortical plate, SP/IZ= subplate/ intermediate zone, SVZ= subventricular zone, ST= striatum, V= lateral ventricle. Scale bar 100µm. **(b, i, p)** Dot plots of migration of green and magenta cells. Each dot represents cumulative mean distance from the top of the cortical plate for all cortical neurons in a coronal brain section. Connected dots indicate measurements were made in the same coronal section. **(c, c’, j, j’, q, q’)** Maximum projection of green cells (c, j, q) and magenta cells (c’, j’, q’) from Double UP, Double UP CIP4, and Double UP FBP17 displayed with inverted contrast to emphasize morphology. Scale bar 50 µm. **(d, k, r)** Representative traces of electroporated neurons from each condition to emphasize morphology. **(e, l, s)** SuperPlots showing comparison of the number of processes from individual green and magenta cells. Each dot represents an individual cell. Triangles represent the mean number of cell processes per brain connected with a line to indicate means are from the same brain. **(f, m, t)** SuperPlots showing comparison of the length of longest processes from individual green and magenta cells. Each dot represents an individual cell. Triangles represent the mean number of cell processes for per brain connected with a line to indicate means are from the same brain. **(g, h, u)** Stacked bar graphs of cell polarity percentages, nonpolar= 0 processes, unipolar= 1 process, bipolar= 2 processes, multipolar= 3+ processes. Error bars show SEM. Paired t-test (two-tailed). (b) 6 brains, mNeon: 424±15.3µm, 693 cells, mScarlet: 427±16.2µm, 554 cells. (e) 5 brains, mNeon: 2.9±0.1 processes, 75 cells, mScarlet: 3.4±0.2 processes, 75 cells. (f) 5 brains, mNeon: 29 ±1.4µm, 72 cells, mScarlet: 27±1.4µm, 73 cells. (g) 5 brains, mNeon: nonpolar 0% of cells mean 0.0±0.0 cells, unipolar 6.7% of cells mean 1.0±0.3 cells, bipolar 39% of cells mean 5.8±1.0 cells, multipolar 55% of cells mean 8.2±0.8 cells, n=75 cells. mScarlet: nonpolar 1.3% of cells mean 0.2±0.2 cells, unipolar 4% of cells mean 0.6±0.4 cells, bipolar 28% of cells mean 4.2±1.5 cells, multipolar 67% of cells mean 10±1.8 cells, 75 cells. (i) 7 brains, Control: 386±27.4µm, 860 cells, CIP4: 407±24.4µm, 873 cells. (l) 7 brains, Control: 3.3±0.3 process, 105 cells, CIP4: 1.0±0.2 process, 105 cells. (m) 7 brains, Control: 25±1.2µm, 104 cells, CIP4: 17±1.6µm, 56 cells. (n) 7 brains, Control: nonpolar 0% of cells mean 0.0±0.0 cells, unipolar 11% of cells mean 1.7±0.6 cells, bipolar 18% of cells mean 2.7±0.9 cells, multipolar 70% of cells mean 11±1.3 cells, 105 cells. CIP4: nonpolar 41% of cells mean 6.2±1.6 cells, unipolar 21% of cells mean 3.1±0.6 cells, bipolar 22% of cells mean 3.3±0.9 cells, multipolar 15% of cells mean 2.3±1.2 of cells, 105 cells. (p) 9 brains, Control: 412±17.5µm, 1308 cells, FBP17: 415 ±18.6µm, 832 cells. (s) 7 brains, Control: 2.8±0.1 processes, 105 cells, FBP17: 2.9±0. processes, 105 cells. (t) 7 brains, Control: 21±1µm, 101 cells, FBP17: 19±1.2µm, 85 cells. (u) brain, Control: nonpolar 3.8% of cells mean 0.6±0.4 cells, unipolar 18% of cells mean 2.7±1.5 cells, bipolar 20% of cells mean 3.0±0.7 cells, multipolar 58% of cells mean 8.7±1.9 cells, 105 cells. FBP17: nonpolar 16% of cells mean 2.4±0.9 cells, unipolar 18% of cells mean 2.7±1.6 cells, bipolar 13% of cells mean 2.0±0.7 cells, multipolar 52% of cells mean 7.8±2.2 cells, 105 cells. All values mean±SEM.

Surprisingly, CIP4 overexpression for two days (E14.5+2) still resulted in a minor, but statistically significant inhibition of migration **(Fig. 3h, i)**. CIP4 overexpressing neurons also had greatly reduced complexity **(Fig. 3j, j’, k)**, with very few processes **(Fig. 3l)**. Most neurons were very spherical with few or no processes. This morphology *in* vivo is consistent with the circular morphology and lack of processes documented in CIP4 overexpressing neurons on a flat cell culture substrate (Saengsawang et al., 2012; Saengsawang et al., 2013; Taylor et al., 2019). Focusing only on cells that did produce processes, we found CIP4 expressing cells still had significantly shorter processes than control cells **(Fig. 3m)**. When grouped by polarity, CIP4 overexpression induced a significantly higher number of nonpolar (process-less) cells and significantly fewer multipolar cells compared to controls **(Fig. 3n)**. Together, the effect of CIP4 overexpression on migration and morphology of radially migrating cortical neurons suggests that CIP4 delays or prevents process initiation, leading to a decrease in migration.

While CIP4 overexpression negatively affected migration at both two and four days after electroporation, FBP17 overexpression at two days after electroporation showed similar migration patterns to controls **(Fig. 3o, p)**. Moreover, FBP17 overexpressing cells did not differ significantly in their morphology compared to controls **(Fig. 3q, q’, r)**, with a similar number of processes, length of longest process and polarity compared to controls **(Fig. 3s-u)**. These results indicate that FBP17 overexpression does not appear to affect morphology or migration after two days of expression post electroporation.

### Knockdown of CIP4, but not FBP17, negatively effects migration by promoting process initiation at E14.5 + 2 days

As mentioned above, since CIP4 protein expression decreases during prenatal cortical development (Saengsawang et al., 2012) and shRNA usually requires several days to effectively inhibit translation, we did not expect to detect a difference in migration two days after electroporation of CIP4 shRNA (E14.5+2). Consistent with the results at four days after electroporation **(Fig. 2b, c)**, the scrambled shRNA resulted in no difference between green and magenta neurons in either migration **(Fig. 4a, b)**, process number **(Fig. 4c-e)** or polarity **(Fig. 4g)** two days post-electroporation. However, we did find a statistically significant difference in the length of the longest process between the control (green) and scrambled shRNA (magenta) neurons **(Fig. 4f)**. This result suggests that the scrambled construct was affecting process outgrowth, which is surprising given that FBP17 shRNA1 **(Fig. 4t)** did not show a difference. Thus, we cannot conclude that CIP4 shRNA1 affected process outgrowth significantly **(Fig. 4m)**. However, since none of the other measurements were significantly different between control and scrambled shRNA, we can confidently compare scrambled shRNA to CIP4 and FBP17 shRNA regarding migration, number of processes and polarity, as outlined below.

**Figure 4.**
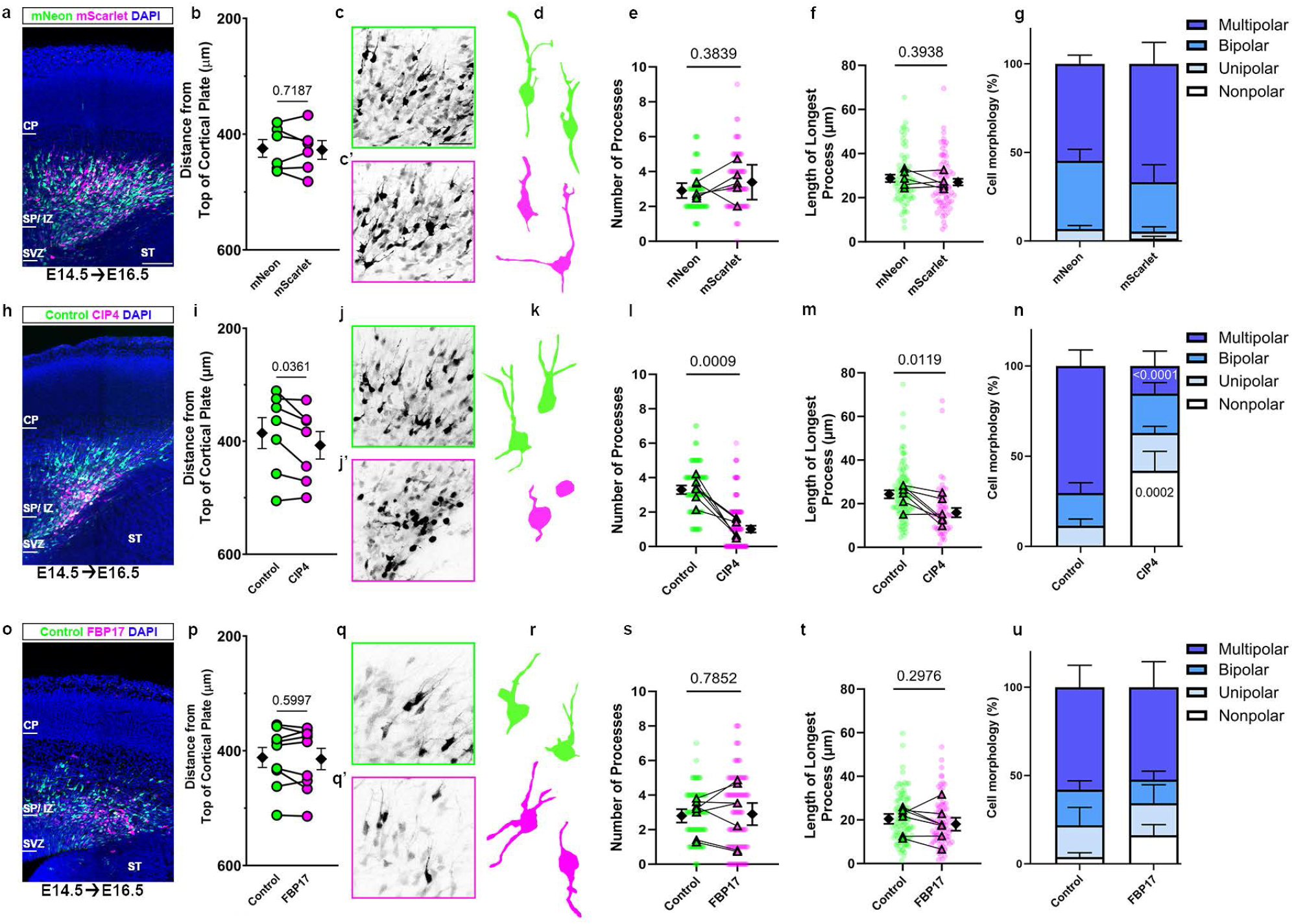
CIP4, but not FBP17, knockdown inhibits migration by decreasing number and length of processes. **(a, h, o)** Representative image of migration after electroporation with Double UP + pSico scramble (a) Double UP + pSico CIP4 shRNA1 (h) and Double UP + pSico FBP17 shRNA1 (o) at E14.5+2. CP= cortical plate, SP/IZ= subplate/ intermediate zone, SVZ= subventricular zone, ST= striatum, V= lateral ventricle. Scale bar 100µm. **(b, i, p)** Dot plot of migration of green and magenta cells. Each dot represents mean distance from the top of the cortical plate to the distance of all cortical neurons in a slice. Connected dots indicate measurements were made in the same coronal slice. **(c, c’, j, j’, q, q’)**. Max project of green (c, j, q) and magenta (c’, j’, q’) cells from Double UP + pSico scramble, Double UP +CIP4 shRNA1, and Double UP + pSico FBP17 shRNA1, displayed with inverted contrast to emphasize morphology. Scale bar=50 µm. **(d, k, r)** Representative traces of electroporated neurons from each condition to emphasize morphology. **(e, l, s)** SuperPlots of number of processes comparing green and magenta cells. Each dot represents an individual cell. Triangles represent the mean number of processes for per condition connected with a line to indicate means come from the same brain. **(f, m, t)** SuperPlots of length of longest processes of green and magenta cells. Each dot represents an individual cell. Triangles represent the mean number of processes for per condition connected with a line to indicate means come from the same brain. **(g, h, u)** Stacked bar graphs of cell polarity percentages, nonpolar= 0 processes, unipolar= 1 process, bipolar= 2 processes, multipolar= 3+ processes. Error bars show SEM. Paired t-test (two-tailed). (b) 6 brains, Control: 412±54.2µm, 887 cells, scramble 410±48.8µm, 586 cells, (e) 5 brains, Control: 2.6±0.2 processes, 75 cells, scramble: 3.0±0.2 processes, 75 cells, (f) 5 brains, Control: 22±1.1µm, 75 cells, scramble: 26±1.3µm, 75 cells. (g) 5 brains, control: nonpolar 0% of cells mean 0.0±0.0 cells, unipolar 27% of cells mean 4.0±2.3 cells, bipolar 25% of cells mean 3.8±0.9 cells, multipolar 48% of cells mean 7.2±2.5 cells, 75 cells. scramble: nonpolar 1.3% of cells mean 0.2±0.2 cells, unipolar 13% of cells mean 2.0±1.3 cells, bipolar 19% of cells mean 2.8±1.0 cells, multipolar 67% of cells mean 10±2.1 cells, n= 75 cells. (i) 6 brains, Control 424±21.2µm, 630 cells, CIP4 shRNA1: 438±22.9µm, 429 cells, (l) 6 brains, Control: 2.9±0.1 processes, 90 cells, CIP4 shRNA1: 4.1±0.2 processes, 90 cells, (m) 6 brains, Control: 19±0.8µm, 88 cells, CIP4 shRNA 1: 27±1.5µm, 89 cells, (n) 6 brains, Control: nonpolar 2.2% of cells mean 0.3±0.2 cells, unipolar 4.4% of cells mean 0.6±0.5 cells, bipolar 31% of cells mean 4.7±0.4 cells, multipolar 62% of cells mean 9.3±0.6 cells, 90 cells. CIP4 shRNA1: nonpolar 0% of cells mean 0.0±0.0 cells, unipolar 1.1% of cells mean 0.2±0.2 cells, bipolar 12% of cells mean 1.8±0.3 cells, multipolar 87% of cells mean 13±0.4 cells, 90 cells. (p) 7 brains, Control: 340±35.4µm, 892 cells, FBP17 shRNA 1: 336±30.4µm, 602 cells, (s) 5 brains, Control: 2.0±0.1 processes, 75 cells, FBP17 shRNA1: 2.1±0.2 process, 75 cells, (t) 5 brains, Control: 19±1.2µm, 70 cells, FBP17 shRNA1: 23±1.3µm, 72 cells, (u) 5 brains, Control: nonpolar 5.3% of cells mean 0.8±0.6 cells, unipolar 31% of cells mean 4.6±0.6 cells, bipolar 39% of cells mean 5.8±0.5 cells, multipolar 25% of cells mean 3.8±1.0 cells, 75 cells. FBP17 shRNA1: nonpolar 4.0% of cells mean 0.6±0.4 cells, unipolar 32% of cells mean 4.8±1.2 cells, bipolar 33% of cells mean 5.0±1.1 cells, multipolar 31% of cells mean 4.6±1.2 cells, 75 cells. All values mean±SEM.

Despite the shorter time for shRNA expression (two days), CIP4 knockdown significantly inhibited migration **(Fig. 4h, i)**, suggesting radial migration is very sensitive to levels of CIP4 expression. Additionally, there was a pronounced effect of CIP4 knockdown on neuronal morphology **(Fig. 4j, j’, k)**, resulting in more processes relative to adjacent, control cells **(Fig. 4l)**. Our data suggests there may be longer process as well, but this cannot be determined with this approach due to the similar difference in process length seen in the scrambled shRNA control condition **(Fig. 4m)**. CIP4 knockdown also resulted in significantly more multipolar cells, and fewer bipolar cells compared to controls **(Fig. 4n)**. Consistent with previous CIP4 knockout data in culture (Saengsawang et al., 2012), we show that CIP4 knockdown resulted in precocious process outgrowth.

We also examined FBP17 knockdown at E14.5+2 and saw no significant effect on migration compared to controls **(Fig. 4o, p)**. Similarly, the morphology of the FBP17 knockdown neurons was not different than controls **(Fig. 4q-u)**. These results suggest that FBP17 overexpression or knockdown has a more nuanced effect on radial migration that is not detected by our morphology measurements at two days post-transfection. We then analyzed the morphology of radially migrating neurons after FBP17 overexpression or knockdown for three days and detected a trend in either more processes (FBP17 overexpression) or fewer processes (FBP17 knockdown) that were not statistically significant (data not shown). Thus, four days of expression of FBP17 or FBP17 shRNA is required to produce migration defects, suggesting that FBP17 may be functioning later in radial migration than CIP4.

### CIP4 expression tapers throughout embryonic cortical development

We have shown previously, via western blot of cortical lysates, that CIP4 expression decreases throughout development and is practically absent in the cortex postnatally through adulthood (Saengsawang et al., 2012). After observing the strong phenotype caused by CIP4 knockdown we wanted to further understand where and when CIP4 protein is present in the embryonic mouse cortex. Utilizing a CIP4 knockout mouse (Hartig et al., 2009), we first validated commercially available CIP4 antibodies via immunohistochemistry and found they all produced a signal indistinguishable from wild-type cortex (data not shown), as we have shown previously in cultured neurons (Saengsawang et al., 2012). We then turned to CRISPR/Cas9 to generate a novel transgenic mouse, in which an mScarlet fluorophore is fused via a flexible linker to the C-terminus of endogenous CIP4. By sectioning the cortices of these CIP4-mScarlet mice and labeling them with an mScarlet antibody, we can for the first time examine expression of CIP4 within intact tissue **(Fig. 5a, a’)**. For comparison, cortices of wildtype littermates were collected, stained for mScarlet, and imaged using identical methods and settings **(Fig. 5b, b’)**. At E12.5, before the cortical plate has been formed, CIP4 expression spans the cortex, from the ventricle to the pial surface, a finding notably absent in the wildtype littermate **(Fig. 5a, b)**. At E14.5, CIP4 expression is absent from the cortical plate and continues to decrease as the cortical plate expands at E16.5 **(Fig. 5a)**. By E16.5 expression is decreased in both the cortical plate and upper intermediate zone, relative to the ventricular and subventricular zone **(Fig. 5a)**. It therefore appears that CIP4 is present in neuronal progenitors and newly born neurons but is depleted or otherwise removed prior to or as neurons enter the cortical plate. This pattern continues at E18.5 with progressive reduction of CIP4 in the cortical plate and intermediate zone **(Fig. 5a)**. As we have shown in previous work with western blots of cortical lysates (Saengsawang et al., 2012), CIP4 expression is essentially undetectable by P10 **(Fig. 5a)**. These data suggest that CIP4 is present in the cortex in a spatial and temporal manner consistent with a role in neuronal migration but appears to not play a role in post-migratory neurons **(Fig. 6)**.

**Figure 5.**
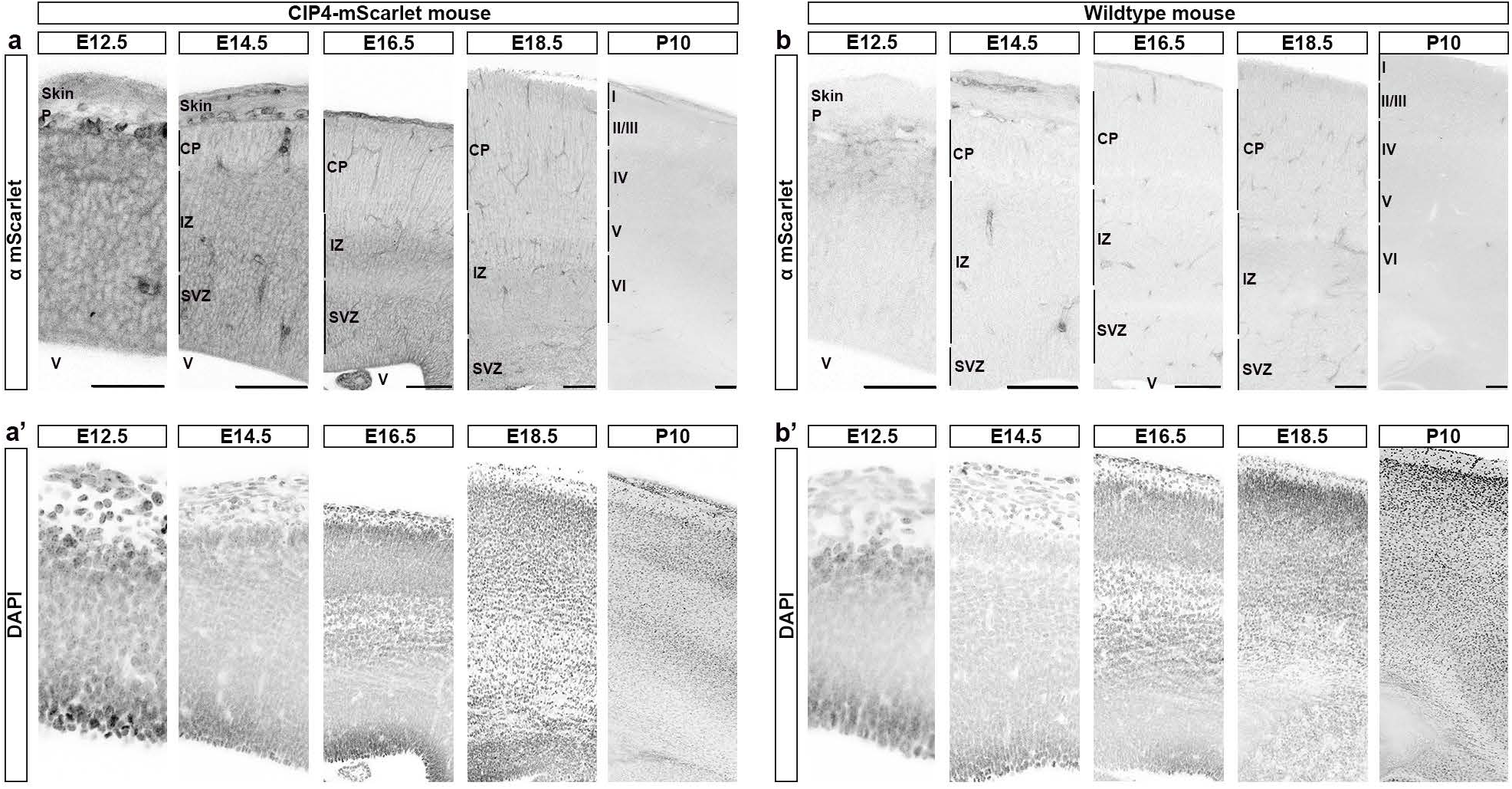
Cortical expression of CIP4 decreases during embryonic development. **a)** Endogenous expression of CIP4-mScarlet in the mouse cortex from E12.5 though P10, stained for mScarlet. **a’**) DAPI staining of corresponding sections from a. **b)** WT litter mates of mice in a, stained for mScarlet expression. **b’)** DAPI staining of corresponding sections from b. All images are shown in inverted contrast to show detail. P= pial surface CP= cortical plate, IZ= intermediate zone, SVZ= subventricular zone, V= lateral ventricle, I= cortical layer 1, II/III= cortical layers 2 and 3, IV= cortical layer 4, V= cortical layer 5, and VI= cortical layer 6. Scale bar 50µm for E12.5 and 100 µm for all other timepoints.

**Figure 6.**
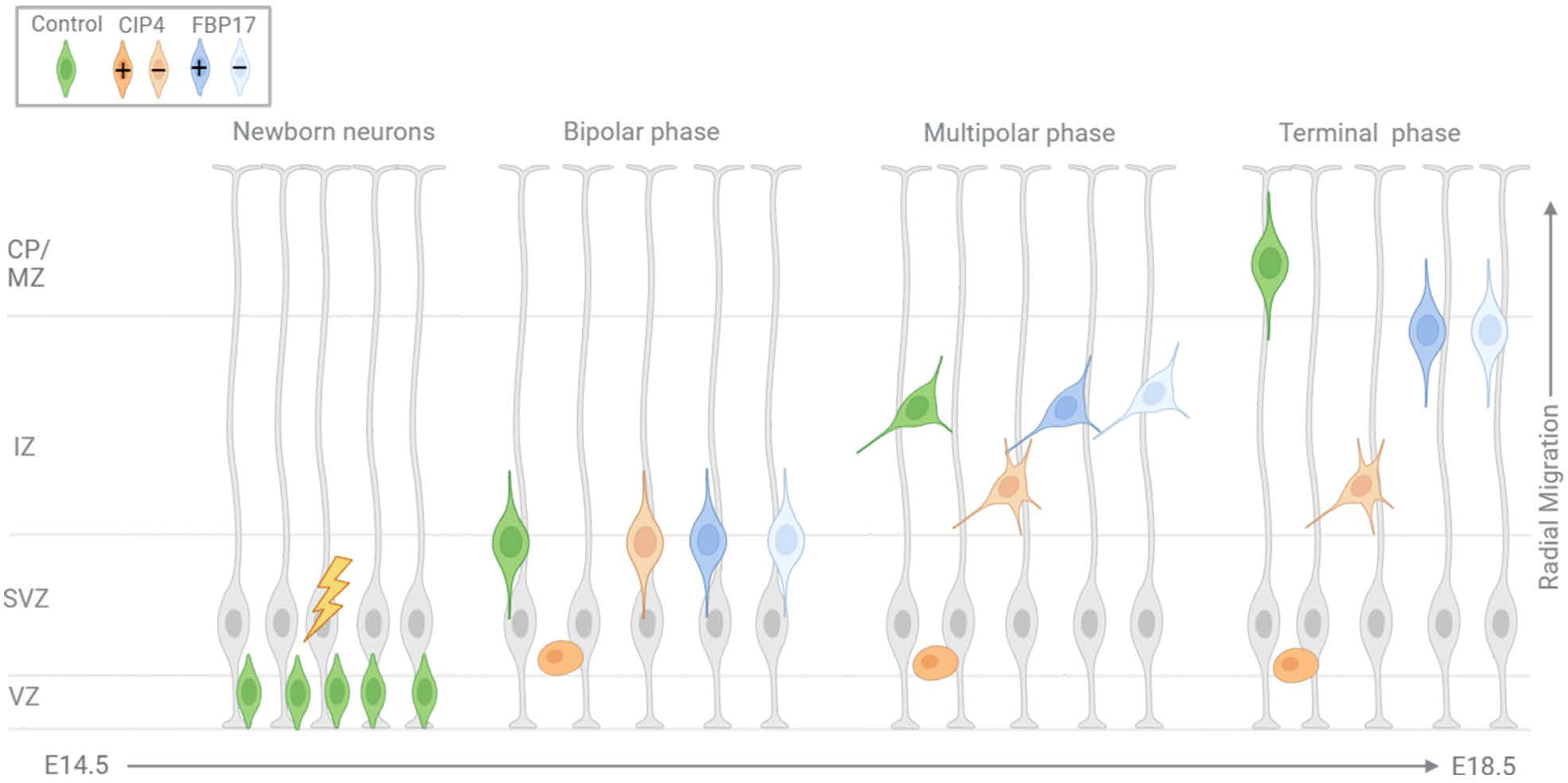
CIP4 and FBP17 have differing effects on radial migration. Normally, a newborn neuron (green) assumes a bipolar morphology soon after birth and migrates from the subventricular zone to the intermediate zone (bipolar phase). In this study we electroporated cells in the ventricular zone at E14.5 (lightning bolt). When the wildtype (green) neuron reaches the intermediate zone the neuron transitions to a multipolar morphology. The, the multipolar neuron transitions back into a bipolar morphology as it enters the cortical plate during the terminal stage of migration. High CIP4 expression (+CIP4) leads to a spherical cell, with few if any processes, that is trapped in the subventricular zone (dark orange) throughout embryonic cortical development. Conversely, CIP4 knockdown (-CIP4) leads to highly multipolar cells trapped in the intermediate zone (light orange). Changes to FBP17 expression (+FBP17, - FBP17) slow migration between E17.5 and E18.5, but do not affect morphology (dark blue and light blue). CP/MZ= cortical plate/marginal zone, IZ= intermediate zone, SVZ= subventricular zone, VZ= ventricular zone. Created in BioRender. English, L. (2024) BioRender.com/o96j986.

## DISCUSSION

Evidence presented here suggests that CIP4 and FBP17 have important roles in brain development by modulating neurite formation and neuronal migration. Premature loss of CIP4 results in more and longer neurites, and prolonged overexpression of CIP4 inhibits neurites almost entirely, resulting in a robust migration deficit. Both loss of FBP17 and overexpression of FBP17 cause a migration defect at E14.5 +4, however, we did not detect a significant difference in the morphology of cells transfected with FBP17 or FBP17 shRNA compared to controls. Thus, we hypothesize that precisely timed CIP4 downregulation is important for the formation of the bipolar morphology that cortical neurons must develop to promote their migration out of the ventricular and subventricular zones, whereas FBP17 plays an important role after the multipolar-to-bipolar transition that occurs later during radial migration **(Fig. 6)**. These findings are consistent with the opposing phenotypes established in our previous cell culture work (Taylor et al., 2019).

The first stage of radial migration occurs in the ventricular zone as the newborn neuron begins process initiation and transitions from a spherical to a bipolar morphology. This transition appears to be important for the subsequent migration of neurons up the radial glia cells toward the subventricular zone (Hatanaka and Murakami, 2002; Noctor et al., 2004). We have shown that CIP4 overexpression inhibits newborn neurons from initiating processes, leading to an early halt in migration, with many cells incapable of migrating out of the subventricular zone **(Figs. 1, 3)**. This phenotype is not due to the death of CIP4 overexpressing neurons. Live imaging of CIP4 overexpressing neurons in organotypic cortical slices showed they oftentimes extended and retracted short processes (data not shown). Moreover, several electroporated litters were allowed to be born and develop to postnatal day 21. In these animals we discovered that at least a portion of CIP4 overexpressing neurons did manage to migrate out of the subventricular zone. However, the neurons only migrated to the intermediate zone and lower cortical plate (data not shown). Processes that extended from these neurons were stunted and extended in random directions, making it difficult to discern if they were axons or dendrites. Unlike control neurons, these CIP4 overexpressing neurons also did not extend through the corpus callosum to the contralateral cortex. These data suggest that CIP4 must be downregulated for neurons to migrate out of the subventricular zone and continued CIP4 expression results in spherical neurons that cannot migrate radially and otherwise progress in the developing cortex.

The second stage of migration occurs in the intermediate zone as neurons transition to a multipolar morphology (Tabata and Nakajima, 2003; Noctor et al., 2004; Namba et al., 2014). When CIP4 is knocked-down, neurons show a decrease in migration as early as E14.5+2 days, which persists to E14.5+4, with many CIP4 knockdown cells unable to translocate to the intermediate zone **(Figs. 2, 4)**. Additionally, there is an increase in multipolar cells among CIP4 knockdown cells compared to controls at E14.5+2 **(Fig. 4)**. We hypothesize that under endogenous conditions CIP4 activity plays an important role in regulating neurite initiation and thus the migratory potential of newborn neurons. Our previous work *in vitro* has established that loss of CIP4 is associated with premature neurite outgrowth, while overexpression of CIP4 causes inhibition of neurite outgrowth (Saengsawang et al., 2012). We believe that the tapering of CIP4 expression in the cortex throughout prenatal development **(Fig. 5)** aids in the neurons developing a multipolar morphology at the correct time and place and allows them to exit the intermediate zone as a bipolar cell capable of migrating into the cortical plate. Thus, the timing and levels of CIP4 expression must be precisely controlled for normal morphology and migration.

The final stage of migration occurs as cells exit the intermediate zone and transition back to a bipolar morphology and translocate through the cortical plate (Noctor et al., 2004). While the purpose of neurons undergoing a multipolar phase is still unclear, failure to exit this phase of migration can result in lissencephaly, epilepsy and intellectual disability (Stouffer et al., 2016). An increasing number of proteins are found to be important for transitioning out of the multipolar stage, including Lis1 (Tsai et al., 2005), doublecortin (Bai et al., 2003), filamin A (Nagano et al., 2004), lamellipodin (Pinheiro et al., 2011) and Nyap1 (Wang et al., 2020). All these proteins are cytoskeletal-associated proteins, suggesting that remodeling of the cytoskeleton is essential for this morphological transition from multipolar to bipolar morphology upon exiting the intermediate zone during migration. Although CIP4 and FBP17 bind actin-associated proteins such as WASP and WAVE, it is thought that their main function is sensing and inducing membrane curvature (Tsujita et al., 2006; Takano et al., 2008; Fricke et al., 2009; Hartig et al., 2009; Saengsawang et al., 2013; Stanishneva-Konovalova et al., 2016). While, two other F-BAR proteins, srGAP2 and GAS7 (Guerrier et al., 2009; Zhang et al., 2016), have been implicated in neuronal migration, CIP4 and FBP17 would be the first F-BAR proteins shown to have a specific role in the multipolar to bipolar morphological transition.

Our data suggests that FBP17 may have a more delayed role in neurite dynamics during radial migration relative to CIP4. Endogenous FBP17 expression begins to increase in the brain after E16.5 (Wakita et al., 2011). Overexpression of FBP17 had an inhibitory effect on migration four days after electroporation, but not at two days **(Figs. 1, 3)**. While it is interesting that early overexpression of FBP17 did not produce an effect on migration or morphology at E14.5 +2, it is possible that the role of FBP17 in neurite dynamics is more subtle, hence harder to detect *in vivo*. This hypothesis is corroborated by our data showing that knockdown of FBP17 does not affect migration until after E14.5 +2, when FBP17 mRNA is likely present. Furthermore, at E14.5 +4 FBP17 overexpressing cells appeared to have significantly longer leading processes compared to controls (data not shown). Considering these findings, we hypothesize that FBP17 is more important for the final stages of migration as cells exit the multipolar stage and enter the cortical plate. Additionally, FBP17 has been shown to play a role in neurite branching and dendritic spine development (Fujita et al., 2002; Wakita et al., 2011) indicating the importance of FBP17 in later developmental stages.

Although CIP4 was originally named Cdc42 interacting protein 4 (Aspenstrom, 1997), in cortical neurons we have shown that CIP4 functions in protruding veils (Saengsawang et al., 2012) and its intracellular distribution to areas of protruding plasma membrane is dependent on Rac- 1, rather than Cdc42 (Saengsawang et al., 2013; Taylor et al., 2019). Contrary to CIP4, FBP17 tubulation in cortical neurons is strongly influenced by Cdc42 activity, but not Rac1 activity (Taylor et al., 2019). Both Rac1 and Cdc42 have been shown to play prominent roles in neuronal migration (Kawauchi et al., 2003; Konno et al., 2005). If during radial migration, CIP4 and FBP17 function with Rac1 and Cdc42, respectively, as we have shown they do in neurite generation in culture, we would expect that phenotypes of CIP4 and FBP17 overexpression could be rescued by dominant negative versions of Rac1 and Cdc42, respectively. Conversely, knockdown of CIP4 and FBP17 could be rescued by constitutively active or wild-type versions of Rac1 and Cdc42, respectively. However, performing these experiments we found no significant difference in radial migration compared to either overexpression or knockdown of CIP4 and FBP17 alone (data not shown).

There are several possible interpretations for our negative results in trying to rescue defects in migration of CIP4 and FBP17 overexpression/knockdown by active/inactive Rac1 and Cdc42 GTPases. We have shown here that titrated levels of CIP4, and to a lesser degree FBP17, is critical for proper radial migration and changing activity with constitutively active or dominant negative Rac1 or Cdc42 may be too blunt of an instrument for precisely regulating CIP4 and FBP17 activity. A more nuanced approach may be to regulate GTPase activity by decreasing Rac1/3 GEF activity by knocking down P-Rex1, or GTPase scaffolds such as POSH. Indeed, P-Rex1 (Yoshizawa et al., 2005; Dimidschstein et al., 2013; Li et al., 2019) and POSH (Yang et al., 2012) have similar migration and morphological phenotypes as CIP4. Another possibility is that during radial migration, CIP4 and FBP17 interact with different GTPases, such as Rnd2 and Rnd3. In the developing cortex Rnd2 and Rnd3 have similar phenotypes and expression patterns to CIP4 and FBP17, respectively (Heng et al., 2008; Pacary et al., 2011). Thus, further work will be needed to define the cohort of proteins interacting with the CIP4 family of F-BAR proteins during cortical neuron radial migration in the developing cortex.

## ACKNOWLEDGMENTS

We thank all members of the Dent lab for their helpful comments and discussions on the manuscript. Funding for this project was provided by NIH NINDS R01-115400 to E.W.D. and the University of Wisconsin Madison Science and Medicine Graduate Research Scholars fellowship to L.A.E. We thank the Office of the Vice Chancellor for Research at UW-Madison and the Wisconsin Alumni Research Foundation or providing funds for the UW2020 Research Initiative and Anjon Audhya for securing the grant that funded the production of the CIP4-mScarlet mouse. We thank the UWCCC Genome Editing and Animal Models Core, supported by P30 CA014520, for use of its services. We thank the University of Wisconsin-Madison Biotechnology Center Animal Models Core (RRID:SCR_024797) and Advanced Genome Editing Laboratory (RRID:SCR_021070) for their contributions in generating genome edited models.

